# The long distance relationship of regional amyloid burden and tau pathology spread

**DOI:** 10.1101/2024.01.15.575698

**Authors:** Merle C. Hoenig, Elena Doering, Gérard N. Bischof, Alexander Drzezga, Thilo van Eimeren, the Alzheimer’s Disease Neuroimaging Initiative

**Affiliations:** Research Center Juelich, Institute for Neuroscience and Medicine II, Molecular Organization of the Brain, Juelich, Germany; University of Cologne, Faculty of Medicine and University Hospital Cologne, Department of Nuclear Medicine, Cologne, Germany; German Center for Neurodegenerative Diseases, Bonn/Cologne, Germany

**Keywords:** functional connectivity, neuropathology, Alzheimer’s disease, staging, pICA, PET, functional connectivity, Alzheimer’s disease

## Abstract

**Introduction:** Consistent with the amyloid-cascade-hypothesis, we tested whether regional amyloid burden is associated with tau pathology increases in spatially independent brain regions and whether functional connectivity serves as a mediator bridging the observed spatial gap between these pathologies.

**Methods:** Data of 98 amyloid-positive and 35 amyloid-negative subjects with baseline amyloid (18F-AV45) and longitudinal tau (18F-AV1451) PET were selected from ADNI. Annual tau change maps were computed. All images were z-transformed using the amyloid-negative subjects as reference. Z-maps of baseline amyloid and annual tau change were submitted to a parallel independent component analysis in GIFT, yielding six component pairs linking spatial patterns of baseline amyloid to longitudinal tau increase. Next, we used the region of maximum coefficient per component as seeds for functional connectivity analyses in a healthy control dataset. This resulted in six pairs of amyloid and tau seed-based networks (SBN). The spatial overlap between these SBNs and components (amyloid OR tau change) and the combined component pairs (amyloid AND tau change) were quantified.

**Results:** Amyloid SBNs presented greater spatial overlap with their respective amyloid components (24%-54%) than tau SBNs with the respective tau change components (16%-40%). However, the spatial combination of amyloid and tau component pairs showed highest spatial overlap with the amyloid SBNs (up to 62% vs. 39% for the tau SBNs).

**Conclusion:** Mechanistically, regional associations of amyloid and tau pathology may be driven by underlying large-scale functional networks. Functional connections may thereby transmit soluble amyloid to remote brain regions within the same network, likely triggering tau aggregation.

## Introduction

Alzheimer’s disease is a dual proteinopathy characterized by the extracellular deposition of amyloid beta (Aß) plaques and the intracellular aggregation of tau neurofibrillary tangles (NFTs). According to the prominent amyloid cascade hypothesis (1), deposition of Aß is the initial event, which triggers the formation of plaques and NFTs, as supported by several in vivo PET imaging studies (2-4). The spatial disconnection between prototypical sites of Aß and tau pathology remains a feature of AD yet unexplained by the amyloid cascade hypothesis.

Histopathological and recent PET imaging studies indicate that these two proteinopathies follow distinct topographies, which initially occur in distal regions of each other. Histopathologically, the distribution of amyloid has been summarized by t Thal phases (5) and the spreading of tau pathology by the Braak stages (6, 7). Recent tau PET imaging studies were able to recapitulate the Braak staging scheme using Flortaucipir (8) and 18F-MK-6240 (9) *in vivo,* but the individual Thal phases could not be easily reproduced using amyloid PET (10, 11). Yet, more recently four distinct stages of amyloid burden were identified based (12). This evidence indicates that PET imaging can be used for staging of both amyloid and tau pathology. Despite the unresolved spatial dissociations observed in these studies, more information on the temporo-spatial interplay of these two pathologies have recently been gathered *in vivo* using PET imaging. Notably, PET imaging currently only captures insoluble forms of Aß and misfolded tau proteins (i.e. NFTs). Nonetheless, using this technique, several studies demonstrated that (antecedent) global Aß plaque load was associated with a greater increase of regional tau tangle pathology (3, 13-15). These studies indicate that Aß plaque deposition may precede neocortical tau aggregation and expedite the build-up and spread of tau pathology. A mechanistic explanation has recently been provided demonstrating that elevated Aß deposition results in higher soluble p-tau concentrations, which was associated with subsequent tau tangle increase assessed with PET (4). Faster tau accumulation rates were in turn observed in regions with strong functional connectivity (4). Regional functional connectivity has consistently been associated with the accumulation of both, Aß plaques (16, 17) and tau pathology (18-20).

While these studies may overall provide some insight into the interplay of global amyloid and regional tau pathology, the spatial disconnect between the prototypical sites of the two pathologies still remains to be deciphered. Importantly, regional amyloid plaque deposition, in contrast to global amyloidosis, may be more specific regarding clinical stage and progression (Pfeil et al. 2022). Therefore, this study was divided into two subparts: First, we aimed to examine the relation between regional amyloid deposition patterns and tau pathology increases using the data-driven approach of parallel independent component analysis (pICA) in a cohort of amyloid-positive subjects. Second, we tested whether the spatial patterns of the component pairs overlap with distinct resting-state connectivity networks, employing a seed-based connectivity approach. We anticipated that the pICA would result in regional components of amyloid burden that were spatially independent of subsequent tau increase patterns, which would fall within the same large-scale functional connectivity network. Presumably this would indicate that regional amyloidosis stimulates tau tangle increase within the same neuronal network.

## Materials & Methods

### Participants

Data for this study was retrieved from the Alzheimer’s Disease Neuroimaging Initiative (ADNI, http://adni.loni.usc.edu/) in May 2022. Key inclusion criteria for the selection of participants were: 1) at least two (baseline and follow-up) evaluable Fortaucipir PET scans not more than three years apart of each other, 2) baseline Florbetapir PET scan, 3) scan interval between the baseline amyloid and tau scan < 6 months, 4) amyloid positivity (mean SUVr > 1.1), 5) ApoE4 carriership information, 6) demographic information available. This resulted in 98 individuals fulfilling the criteria in ADNI, comprising cognitively-unimpaired (CU) and 50 cognitively impaired subjects (Table 1). In addition, a reference group of amyloid-negative CU subjects was included, for whom longitudinal tau-PET scans and a baseline amyloid scan was available (Table 1). All procedures performed in this study were in accordance with the ethical standards of the institutional and/or national research committee.

**Table 1.** Demographic characteristics of the groups included in this study. Mean and standard deviations are provided for continuous variables. CU = cognitively unimpaired; CI = cognitively impaired, Amy -/+ = amyloid negative/positive based on SUVr of 1.1; SUVr = standard uptake value ratio. P-values are based on the Kruskal-Wallis (across group) or Mann-Whitney-U (Amy+ CU vs. Amy+ CI) tests for continuous and Chi squared (across groups) and Fisher exact (Amy+ CU vs. Amy+ CI) test for categorical variables. * For one participant in the amyloid negative CU group, the ApoE4 information was unavailable.

| Demographics | Amy- CU<br>N=35 | Amy+ CU<br>N=48 | Amy+ CI<br>N=50 | Group differences |  |
| --- | --- | --- | --- | --- | --- |
|  |  |  |  | Across<br>groups | Amy+ CU<br>vs. Amy+ CI |
| Age | 74.66 (8.09) | 75.01 (6.07) | 75.78 (6.67) | p=.104 | p=.486 |
| Sex (M/F) | 16/19 | 15/33 | 27/23 | p=.073 | p=.026* |
| ApoE4 (pos/neg) | 8/26* | 25/23 | 32/18 | p < .001* | p = .306 |
| ADAS 13 | 6.82 (3.64) | 8.78 (5.47) | 20.20 (10.69) | p<.001* | p<.001* |
| Education (years) | 15.74 (2.79) | 16.98 (2.04) | 16.00 (2.70) | p=.068 | p=.057 |
| Global Amyloid (SUVR) | 1.01 (.04) | 1.33 (.20) | 1.40 (.16) | p < .001* | p = .019* |
| Scan Interval (years) | 1.53 (.62) | 1.32 (.54) | 1.31 (.54) | p=.438 | p=.994 |

### PET image processing

Flortaucipir (tau) and Florbetapir (amyloid) PET scans were co-registered to the structural MRI, normalized to the tissue probability map, and subsequently smoothed with a Gaussian kernel of 6mm FWHM using SPM12 (Wellcome Trust Centre for Neuroimaging, London, UK). Standard uptake value ratios (SUVRs) images were computed using the inferior cerebellar grey matter as reference region for the tau scans (21, 22) and the whole cerebellum for the amyloid scans (23). Next, amyloid PET scans were z-transformed using the baseline amyloid scans of the amyloid negative control group. The respective tau PET scans (baseline and follow-up) were z-transformed using the mean of the baseline and follow-up tau PET scans of the amyloid negative control group. Finally, annual tau increase maps were computed (T1-BL/Scan Interval). The pre-processed z-maps were then submitted to further analysis.

### Parallel independent component analysis

We previously employed independent component analysis on tau PET data to identify regional components of baseline tau pathology (20). Since, in the current study, we aimed to link the spatial information of amyloid and annual tau change, we employed the data-driven approach of a parallel independent component analysis (pICA). Briefly, the goal of pICA is to identify weighted combinations of two modalities that are linked with each other. It is a multimodal analysis that, in a first step, extracts components that are maximally independent within each of the two modalities and in second step it finds the interconnection between the two component sets of the different modalities or PET tracers by maximizing the linkage function in a joint estimation process, through canonical correlation (24).

Here, we entered the baseline amyloid z-maps as one feature and the annual tau change maps as second feature to the pICA. The analysis was conducted using the Fusion toolbox in GIFT (https://trendscenter.org/software/fit/). In a first step, dimensionality reduction was performed by principal component analysis. The resulting principal components were afterwards submitted to the pICA. The infomax algorithm was employed to identify independent components in the amyloid and annual tau change patterns. The component pair number was set to a maximum of eight, which was estimated by the minimum description length criteria. The resulting component pairs were afterwards assessed in terms of plausibility, leading to the exclusion of one amyloid and one tau component given that they clearly represented artefact patterns. This resulted in six remaining component pairs, which were thresholded at z > 1.96 and finally, the mean z-scores of baseline amyloid burden and annual tau increase were extracted for the respective IC maps for each individual.

### Seed-based functional connectivity analysis

To link the component pairs by a potential mediating mechanism, we conducted a seed-based connectivity analysis using pre-processed resting-state data of a healthy control dataset from the 1000 functional connectome project (http://fcon_1000.projects.nitrc.org/; (25)). The dataset included 26 healthy controls (13 males/13 females) aged between 23 and 44 years (mean= 29.77 ± 5.21 years). More detailed information on the pre-processing of the data can be found elsewhere (20). Briefly, the resting-state data was analysed using a common pipeline of the DPARSF toolbox (26). As seeds for the functional connectivity analysis, we extracted the coordinate of the maximal z-score from each of the six amyloid and five tau components. One of the tau components was correlated with two individual amyloid components. The seeds were then defined as a sphere centred at the respective peak coordinate with a radius of 6 mm. Afterwards, the correlation matrix of the resting-state functional MRI timeseries was computed between the respective seed and coherently active regions yielding six amyloid and five tau functional connectivity maps for each individual. In a final step, we conducted one-sample t-tests in SPM12 (FWE corrected, p <.05) with the individual functional connectivity maps, which resulted in the respective amyloid or tau seed-based networks (SBN). The SBNs were then binarized using the t-score threshold that was based on the FWE-correction.

### Statistical analyses

#### Spatial characterization of component pairs according to staging schemes

To characterize the spatial pattern of the respective component pairs, the binarized maps of the amyloid baseline and tau change components were overlayed with either the previously established Grothe stages (12) or the Braak stages (as previously employed for Flortaucipir, (27)). The spatial overlap between the components and the respective staging schemes was quantified using Dice Similarity Coefficient (DSC), which yields the percentage spatial overlap between two binary volumes:

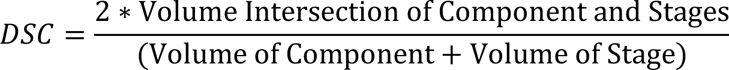

The highest DSC score was chosen as the best fit for the respective components.

Moreover, we also computed the spatial overlap between the component pairs, themselves, using the DSC score. The DSC score is interpreted as follows: <0.2 reflects poor, 0.2–0.4 fair, 0.4–0.6 moderate, 0.6–0.8 good, and >0.8 near complete overlap (28).

#### Functional connectivity and spatial extension of regional amyloid and tau pathology

To examine the potential role of functional connectivity as mediator bridging the spatial gap between regional amyloid and tau pathology, we further quantified the spatial overlap between the SBNs derived from the amyloid components with the independent components (amyloid baseline OR annual tau change component) as well as with the combined component pair (spatial combination of the amyloid AND the annual tau change component of the respective pair). Notably, this time we did not use the DSC as the DSC score considers the sum of both volumes in the denominator, which would consequently change for each comparison. Since we were interested in the added value of the component pairs, we computed the relative spatial overlap with the respective SBN the following:

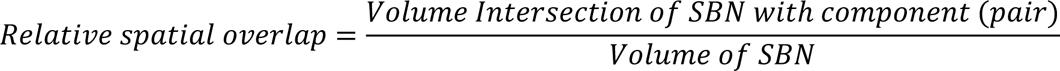

Thus, the resulting value provided information on which percentage of the network was covered by the respective component or component pairs.

#### Correlation and regression analyses

As a proof of concept, we performed spearman analyses between the extracted baseline amyloid z-score and the tau change z-score of each component pair correcting for sex, age, ApoE4 and MMSE score.

Next, to assess group-specific effects of the component pairs, linear regression modelling was used for each group separately examining the predictive power of mean amyloid load within the respective component, ApoE4 carriership, sex, and age on the annual increase of tau pathology in the corresponding tau component. Bonferroni correction acccounted for multiple comparisons (0.05/12=.004).

For all conducted regression models, the mean tau z-scores were logarithmically transformed to fulfill the assumptions of normal distribution of residuals and homoscedasticity. Extreme residual outliers with a standard deviation < -2 or > 2 were excluded from the respective models.

All analyses were performed in SPSS 28. Graphical illustration was performed using R Studio and MRIcron.

## Results

### Group characteristics

The amyloid-negative CU group significantly differed by design in terms of global amyloid load *(H(2)=79.77, p <.001)*, ADAS13 *(H(2)=55.71, p <.001)* and presented a lower frequency of ApoE4 carriers (χ*2=13.506, p = .001)*. Age, sex, education and scan interval did not differ across groups. The amyloid positive CU group differed from the amyloid-positive CI group in terms of sex *(p=.026),* global amyloid load *(U= 1529, Z=2.338, p=.019)* and by design in ADAS13 score *(U=2059, Z=6.106, p< .001)*. Age, education, ApoE4 status, and scan interval did not significantly differ between the two groups. For the statistical summary see Table 1.

### Spatial patterns of regional amyloid and annual tau increase

The pICA resulted in six component pairs (Fig. 1), some of which were characterized by a close spatial overlap (Pairs 1, 3 and 6), but others by a spatial disconnect (Pair 2, 4, and 5). The DSC score ranged from 3% overlap up to 52% between the component pairs.

**Figure 1.**
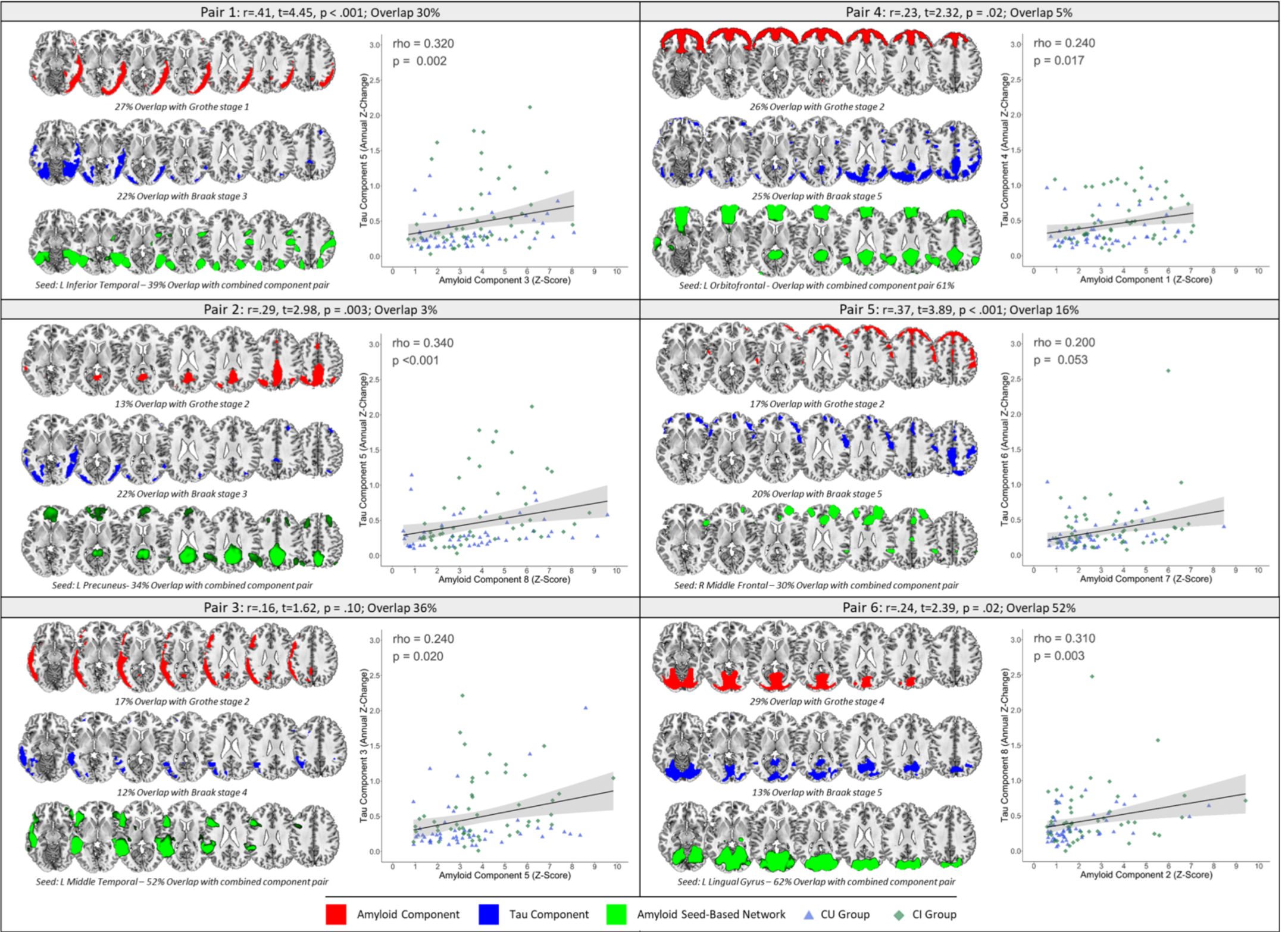
Regional associations of amyloid pathology with annual increases in tau pathology. The six component pairs of the parallel independent component analysis are depicted including the correlation coefficient of the components’ mixing coefficients as well as the spatial overlap, DICE, between the component pairs. The amyloid components are depicted in red and the corresponding tau component in blue. Below the independent components the respective stage according to previously established staging schemes is listed in concordance with the relative spatial overlap. The scatterplots illustrate the association between pathological burden within the component pairs. Additionally, the seed-based network based on the maximum region of the amyloid independent component is shown in green. The spatial overlap between the seed-based network and the combined component pattern of the amyloid and tau impendent components is provided.

Spatial characterization of the component pairs according to previously established staging schemes of amyloid (Grothe stages) and tau (Braak stages) pathology yielded that early amyloid stages (Grothe 1-2) were related to moderate/advanced Braak stages (Braak III-V), whereby the late amyloid stage (Grothe 4) corresponded to Braak stage V. The earliest Braak stages I-II were not captured by any of the tau components, potentially due to primary age-related tauopathy in the control group influencing the z-map deviation maps. The best overlapping stages for each component is listed below the corresponding component in Fig. 1.

Mean amyloid load of the respective amyloid components was significantly associated with increases in mean annual tau burden in the corresponding tau components across groups (with a correlation coefficient of around rho=.25-.35), except in component pair 5, which only reached marginal significance (rho=.200, p=.053). Statistical results are provided in the scatterplots of Fig. 1.

### Functional connectivity and regional associations

The amyloid-derived SBNs presented moderate-to-good spatial overlap with the corresponding amyloid component (24%-54%), but lower spatial overlap with the corresponding tau component (5%-34%, Wilcoxon-test: *z* = -2.201, *p* = .028). Moreover, a non-parametric Friedman test yielded that the spatial combination of the amyloid and tau component pairs showed highest spatial overlap with the corresponding amyloid SBN (up to 62%) in comparison to the amyloid or tau component, separately (χ2(3) =12, p=.002). In contrast, the tau-derived SBNs presented low-to-moderate overlap with the corresponding tau component (16%-40%) and significantly lower or absent overlap with the respective amyloid components (0%-19%; Wilcoxon-test: *z* = -1.992, *p* = .046). When comparing the combination of the amyloid and tau component pairs, the spatial overlap with the respective tau SBNs was significantly lower (25%-40%) compared to the amyloid SBNs (30%-62%; Wilcoxon-test: *z* = -2.009, *p* = .041). The respective percentages of spatial overlap are listed in Table 2.

**Table 2.**
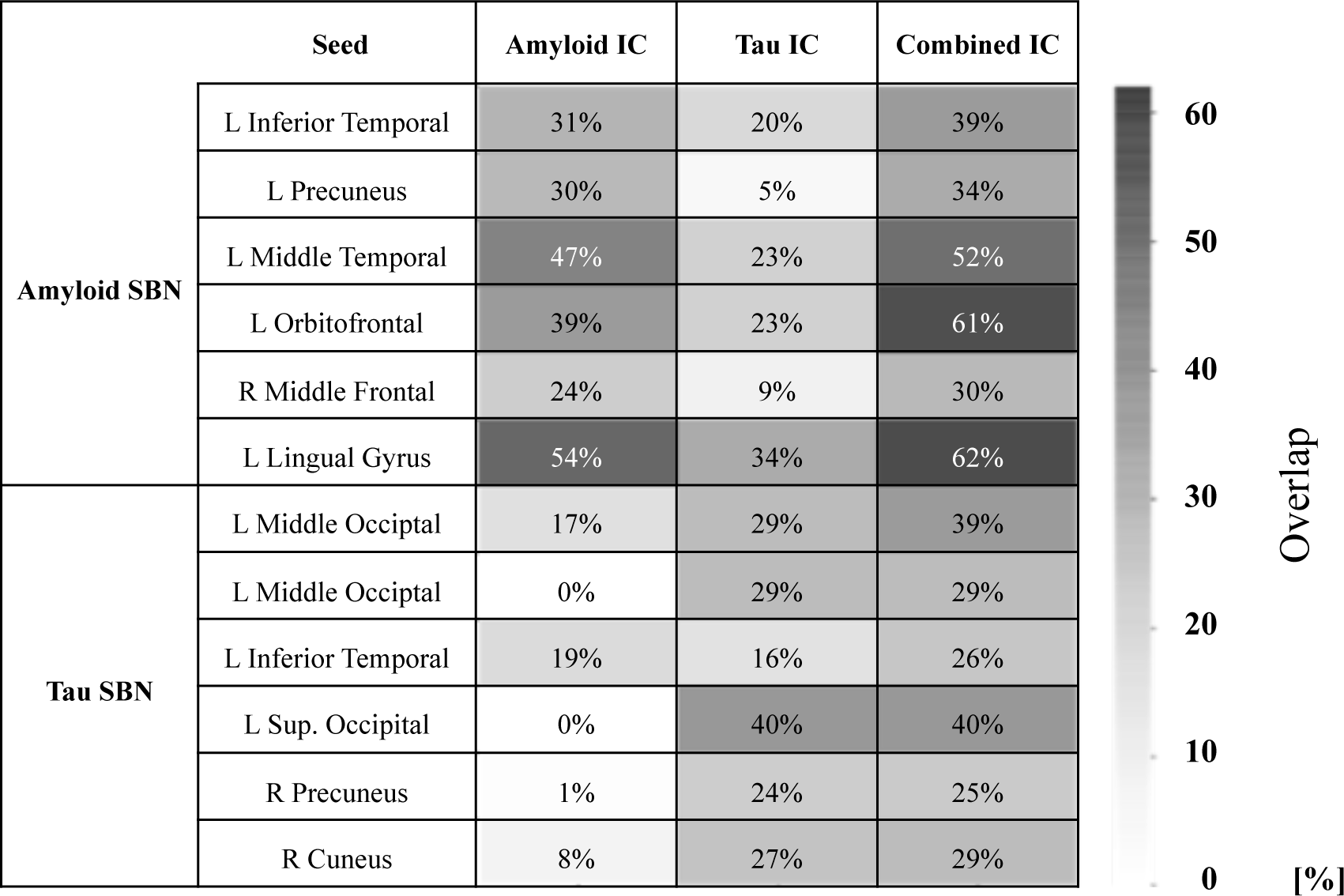
Spatial overlap of the seed-based networks with the identified component patterns. The percentage spatial overlap between the components and the respective SBNs as well as the respective seeds for each component are listed. The seeds in the column next to the amyloid SBN refer to the maximum z-score within the respective amyloid IC of each pair, whereas the seeds in the column next to the tau SBN refer to the maximum z-score region within the respective tau IC. The order of the SBNs corresponds to the numbering in Fig. 1. IC = independent component; SBN= seed-based network.

### Regional associations of amyloid and annual tau increase is dependent on clinical stage

The regression analyses performed for the clinical groups, separately, yielded that amyloid load in component pair 1 (ß=.528, p=.001) and in component pair 5 (ß=.508, p=.001) was the only significant predictor of tau increase in the CU group and not in the CI group (after multiple comparison correction). ApoE4, age, sex were not significant predictors in this model. For the CI group, we found that amyloid load of component pair 3 (ß=.402, p=.003) was a significant predictor of tau increase in the corresponding tau component, but not in the CU group. The regression model for component pair 2 yielded a significant main effect of amyloid load the corresponding amyloid IC in both groups. Interestingly, a significant main effect of ApoE4 carriership was observed for the CI group containing component pairs 1,2,5 and 6. The main effects of sex and age were not significant in any of the models. A summary of the statistics can be found in Table 3.

**Table 3.**
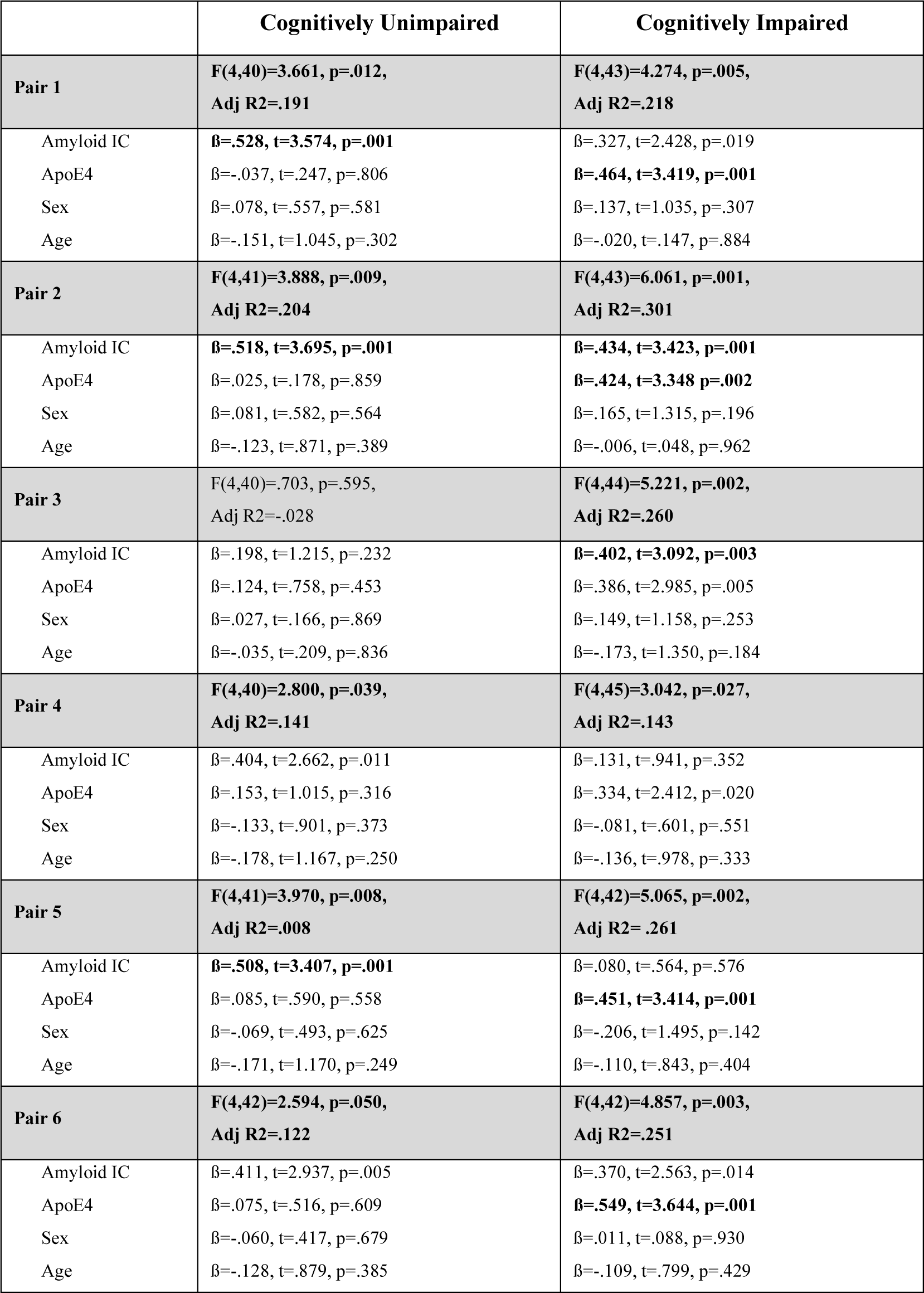
Group-specific effects of regional amyloid on annual increases in tau pathology. The models include the four independent variables (amyloid IC, ApoE4, sex and age) used for predicting the annual increase in tau pathology of the corresponding tau IC (dependent variable). The pairs refer to the component patterns that are illustrated in Fig. 1. The model summary is provided for each group, separately, including the main effects of each predictor, standardized beta coefficient, t-value and significance level. Significant models and main effects (after Bonferroni correction p < .004) are highlighted in bold.

## Discussion

A substantial body of evidence suggests that the adverse effects of Aβ pathology depend on tau, thereby placing Aβ upstream of tau in the pathogenesis of AD (29-31). Here, we identified a set of component pairs of baseline regional amyloid deposition and subsequent tau pathology increases using PET imaging. Several pairs presented a regional co-localization of amyloid and subsequent tau pathology increases (i.e., high spatial overlap), while others were characterized by a spatial discordance (i.e., low or no overlap). Importantly, the identified components roughly followed previously established staging schemes and were specific for the clinical stage, i.e., cognitively unimpaired vs. impaired status. Finally, we observed that the spatial patterns of the component pairs aligned relatively well with functional connectivity networks that were derived from the maximum amyloid-deposition peak within the amyloid components. These results indicate that Aβ-mediated effects on tau pathology aggregation and subsequent spreading occur within the same large-scale cortical networks.

### Diagnostic and prognostic utility of regional patterns of regional amyloid and tau pathology increase

The two characteristic hallmarks of AD evolve in a temporo-spatial order including a long preclinical phase, different spatial affinities (i.e. stages) and consequences on cognitive function (32). By employing the data-driven approach of pICA in a cohort of cognitively (un-)impaired amyloid-positive subjects, a set of regionally-related components of baseline amyloid and subsequent increases of tau pathology were identified among which the neuropathological burden of the two proteinopathies was highly correlated. Interestingly, the spatial characterization of the components according to previously established staging schemes yielded that early amyloid phases (Grothe stages) accorded with moderate Braak stages, whereas the highest amyloid stage was linked to advanced Braak stages arguing that PET imaging can potentially be used in the staging of both amyloid and tau phases. In accordance with the observation that these stages are linked with clinical severity (8, 12, 14, 33), we found that some of the component pairs were specific for clinical stage pointing towards their potential prognostic utility. Indeed, the component pair characterized by an amyloid component comprising orbitofrontal regions and subsequent increases in tau pathology in medio-temporal regions was predominantly expressed by the cognitively unimpaired amyloid-positive group. These results are in line with disease staging suggesting early amyloid phases are linked to predominant orbitofrontal involvement, while early tau pathology relates to medio-temporal vulnerability. Interestingly, the component pair characterized by asymmetric temporal amyloid and spatially overlapping increases of tau pathology was specific for the cognitively impaired group. Supporting this finding, asymmetric patterns of temporal and precuneal amyloid burden have recently been suggested to be predictive of the progression from cognitively normal to impaired stages (34). Accordingly, it has been proposed that local rather than global levels of amyloid burden may be more sufficient to inform on differential disease-trajectories (12, 34). Moreover, it has been suggested that the extent of Aβ plaque burden in specific regions could be employed in the prediction of disease progression (35).

Importantly, the spatial characteristics of the identified component pairs suggest that multi-tracer PET imaging in combination with data-driven approaches such as pICA may permit regional staging resembling neuropathological observations and may provide diagnostic and prognostic clinical value. Notably, the determined component pairs aligned with previously established components of amyloid and tau pathology assessed in a cross-sectional approach using PET imaging (36). This suggests that some of the spatial interrelations already exist at baseline and continue to progress longitudinally across distinct mechanistic pathways, which will be elaborated on the following.

### Functional connectivity bridging the gap between regional amyloid and tau pathology

Several mechanisms have been considered explaining the stereotypical spread of the two proteinopathies of AD. In particular, multimodal imaging studies consistently reported that the topographies of neurodegenerative disease pathologies overlap with large-scale neuronal networks arguing that functional and structural connectivity between regions promotes the distribution of these pathologies (37, 38). Here, we observed a good spatial correspondence of the amyloid spatial patterns with functional connectivity networks that were derived based on the seed regions of the respective amyloid component. More importantly, when assessing the spatial overlap of the corresponding tau components with the amyloid networks, we found that these patterns partly coincided within the same functional network. In contrast, the spatial correspondence of the amyloid components with the tau seed-derived functional networks was poor or even absent. This suggests that the amyloid functional network may be affected upstream and that both, Aβ plaque deposition and tau tangles occur within the affected network. Presumably, transmission of soluble Aβ and the subsequent accumulation of Aβ plaques across this network facilitates the aggregation of tau tangle pathology within distinct regions of the same network. The regional aggregation of tau tangles may then trigger spreading of tau pathology outside the initially affected network, into regions not primarily affected by Aβ plaques.

Following this line of argumentation, evidence based on *in vivo* studies in mice suggests that seed-induced soluble Aβ aggregates usually accumulate around the site of injection, but eventually spread along axonally-connected regions to remote locations of the brain pointing towards structural but also extracellular routes contributing to the spreading process (for review see (39)). Also *in vivo* functional MRI studies reported higher accumulation of insoluble Aß plaques in hub regions of large-scale networks, suggesting that regions with high connectivity and topographical relevance for a given network may promote the transmission of soluble Aβ along the connected pathways (16, 17).

Intriguingly, it has recently been shown that Aß plaque load, as measured with PET, was related with an increase in soluble plasma (4) and CSF p-tau levels (14). Based on these recent findings, expansion of soluble Aß along distinct network pathways might result in an increase in soluble p-tau levels across the affected network, which in turn may eventually lead to the aggregation of insoluble tau tangles in vulnerable regions of the brain. Indeed, Aβ-mediated increase in plasma p-tau was associated with the subsequent accumulation of neocortical tau pathology over time (4). More importantly, higher soluble p-tau concentrations were linked with faster accumulation rates of tau pathology in regions with strong functional connectivity, again pointing towards the notion of functional connectivity being a facilitator in the neuropathological aggregation and spread (4). Given this new in vivo evidence, Aβ plaques may mediate local production of soluble p-tau, which in turn travels to remote brain regions following pathways of strongest connectivity and greater regional vulnerability proteinopathies (40, 41). Eventually, these soluble forms of tau pathology aggregate into insoluble forms within the given network and induce a seeding process of misfolded tau aggregates outside the initially affected network (Fig. 2).

**Figure 2.**
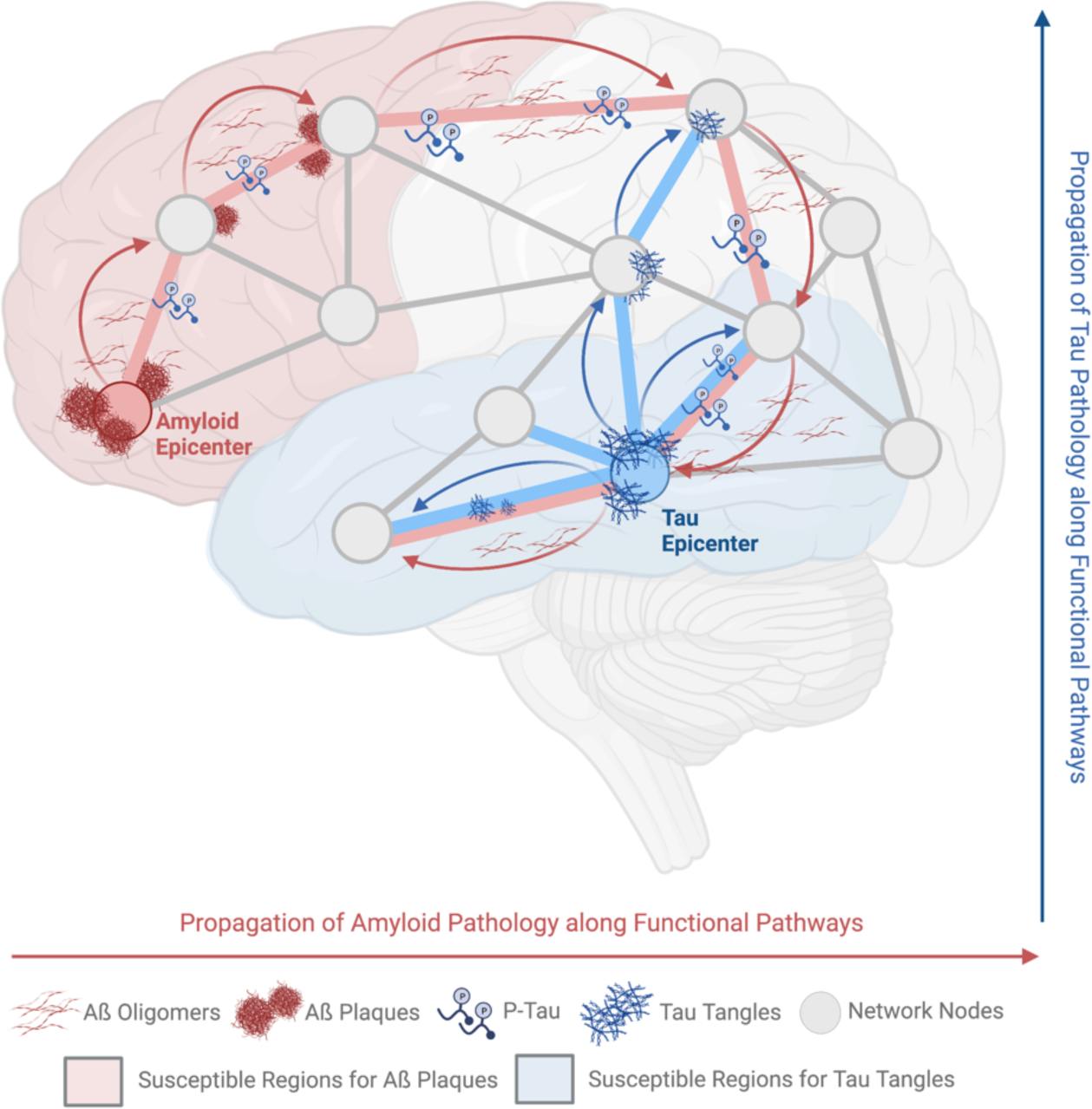
Schematic presentation of the potential link between regional amyloid burden and increase of tau pathology across functional networks. Insoluble amyloid plaques predominantly accumulate in vulnerable regions (red), for which a neurogenetic contribution has been reported (40, 41). Soluble forms of these proteins likely follow pathways of high functional connectivity (i.e. pathways in red). Across these pathways, the presence of insoluble and soluble forms of amyloid may result in an upregulation of soluble phosphorylated tau forms (4). In turn, soluble phosphorylated tau species likely aggregate into insoluble tau tangles in regions (blue) that have been shown to be highly vulnerable to tau pathology (40, 41). Misfolded tau aggregates then cause a seeding process within the same, but eventually also outside the initially affected network (i.e. pathways in blue). The figure was created using biorender.com

Regional vulnerability in combination with the expansion of proteinopathies along functional pathways may cause the observed spatial disconnect between the two proteinopathies. Yet, down-the-line of this pathological cascade, the commonality of these regions, namely the underlying functional connectivity, may finally result in the relative co-localization of these two proteinopathies. Indeed, recent evidence indicated that Aβ asserts its adverse effects as soon as it co-localizes with misfolded tau aggregates, an event which then causes the spreading of misfolded tau proteins into regions that may not primarily be affected by Aβ (42-44). Also in this study, we identified several component pairs with high spatial overlap, which in turn were associated with later disease stages.

The current work supports the growing evidence that the two characteristic hallmarks of AD are strongly tied to structural (45) and functional (18, 20, 21) network architecture. Importantly, network connectivity is likely a facilitating mechanism in the stereotypical distribution of these proteins (as soluble forms) and may cause the eventual co-localization and assembly of these proteinopathies (into insoluble forms) in vulnerable brain regions. Additional independent factors, such as genetic predisposition may further amplify AD pathogenesis such as in case of ApoE4 carriership, as also signified by this study.

A few limitations need to be mentioned when considering the findings of this study. First, we used a group of amyloid-negative subjects as reference to standardize the amyloid and tau PET scans. This group was slightly younger than the amyloid-positive CU and CI group. Yet, it Moreover, the analysis was based on a dataset of younger healthy controls and may thus not encompass the actual underlying network structures of the included patients. However, given that protein aggregation may actually cause network changes, assessment of premorbid (healthy) network structures may provide valuable information on the mechanistic pathways of neuropathology spread (46). Finally, the functional connectivity analysis was limited to the seed of maximum z-score of the amyloid or tau component, respectively. Future studies may thus employ a more refined approach using graph theory using younger and age-matched samples.

Overall, we demonstrated that regional amyloid pathology may inform on subsequent tau pathology increases in spatially overlapping, but also remote brain regions. Importantly, the identified component pairs of regional amyloid and tau tangle increase were associated with clinical stages, indicating that they may provide clinical prognostic utility across the AD spectrum. From a mechanistic point of view, it appears that the regional associations of amyloid and tau pathology increases are facilitated by functional connections within large-scale neuronal networks.

## Acknowledgements

MH received funding from the Alzheimer Forschung Initiative e.V. In addition, this study was supported by the German Research Foundation (DFG, DR 445/9-1). GNB is funded by the Deutsche Forschungsgemeinschaft - Project-ID 431549029 - SFB 1451.

Data collection and sharing for this project was funded by the Alzheimer’s Disease Neuroimaging Initiative (ADNI) (National Institutes of Health Grant U01 AG024904) and DOD ADNI (Department of Defense award number W81XWH-12-2-0012).

